# Binding strength and hydrogen bond numbers between Covid-19 RBD and HVR of antibody

**DOI:** 10.1101/2020.12.21.423787

**Authors:** Ryan Taoran Wang, Alex Fan Xu, Qi Zhou, Tinglu Song, Kelvin J. Xu, Gu Xu

**Author notes:** First author (Ryan Taoran Wang, Alex Fan Xu).

## Abstract

The global battle against the Covid-19 pandemic relies strongly on the human defence of antibody, which is assumed to bind the antigen’s Receptor Binding Domain with its Hypervariable Region. Due to the similarity to other viruses such as SARS, however, our understanding of the antibody-virus interaction has been largely limited to the genomic sequencing, which poses serious challenges to the containment, vaccine exploration and rapid serum testing. Based on the physical/chemical nature of the interaction, infrared spectroscopy was employed to reveal the binding disparity, when unusual temperature dependence was discovered from the 1550cm^-1^ absorption band, attributed to the hydrogen bonds by carboxyl/amino groups, binding the SARS-CoV-2 spike protein and closely resembled SARS-CoV-2 or SARS-CoV-1 antibodies. The infrared absorption intensity, associated with the number of hydrogen bonds, was found to increase sharply between 27°C and 31°C, with the relative absorbance matches at 37°C the hydrogen bonding numbers of the two antibody types (19 vs 12). Meanwhile the ratio of bonds at 27°C, calculated by thermodynamic exponentials rather than by the layman’s guess, produces at least 5% inaccuracy. As a result, the specificity of the SARS-CoV-2 antibody will be more conclusive beyond 31°C, instead of at the usual room temperature of 20°C - 25°C, when the vaccine research and antibody diagnosis would likely be undermined. Beyond genomic sequencing, the temperature dependence, as well as the bond number match at 37°C between relative absorbance and the hydrogen bonding numbers of the two antibody types, are not only of clinical significance in particular, but also of a sample for the physical/chemical understanding of the vaccine-antibody interactions in general.

## Introduction

The ongoing battle against the coronavirus (Covid-19, or SARS-CoV-2), has largely been defined by the human antibody, including the vaccine development, and rapid diagnosis of serum, all based on the match and binding between its Hypervariable Region (HVR) and the antigen’s Receptor Binding Domain (RBD)^[1–7]^. However, the existing knowledge has been confined by the conflicting observations beyond genomic sequencing, such as the early disappearance of the antibody, which poses serious challenges to not only vaccine development^[8,9]^, but also the current state-of-the-art Covid-19 IgM/IgG rapid serum tests, due in part to the similarity to other viruses, such as SARS, and even that of a common flu^[10,11]^.As a result, the progress of vaccine development has been largely hindered and the performance of various antibody tests was undesirable. Studies showed that even vaccines of short term immunity would not be available for another 12-18 months^[12–14]^, and the accuracy of various antibody tests were found to range from 62% to 95%^[15,16]^. All this indicated that the pandemic might become seasonal, when lockdown failed by deficient appraisal, and the longing vaccine exploitation and desirable rapid serum testing deemed unreliable.

To unravel the cause of the conflicting observations relevant to the RBD and HVR binding, it is necessary to first differentiate the binding variation between the SARS-CoV-2 and other less distinguishable coronaviruses, such as SARS-CoV-1, especially in terms of the chemical bonding numbers between the viruses and their respective antibodies^[3,17]^. In particular, one has to clarify, how the binding sites of the spike proteins (S proteins), which are the foremost weapon head of the virus, interact with the antibodies of SARS-CoV-2 (antibody 2), the frontier human defence^[3,18,19]^. Although a number of characterization methods have already been employed to analyze the structure of the coronaviruses, especially their S proteins^[20–22]^, when it was found that the RNA sequence of SARS-CoV-2 bears 89.1% resemblance to the SARS-CoV-1^[21]^, which only deciphers the similar 3D structures of the S proteins between the two viruses obtainable from Cryo-EM^[20]^. It does not, however, provide the details of the binding disparity between the viruses and their antibodies. Neither could the latter be elucidated by the amino acid sequencing of the S proteins^[3,19]^, despite the finding that 77% of the amino acid sequences were identical in the RBD. Consequently, the attempt thus far has been destined to be less than successful, due not least to the lack of probing of the binding difference between the viruses and their antibodies. More precisely, even the bonding nature between the two has yet to be illustrated, let alone the influence of other factors, such as temperature, or the possible antibody dependent enhancement.

It is therefore the purpose of the current letter, to probe the binding strength variation between the SARS-CoV-2 spike protein/SARS-CoV-2 antibody, and SARS-CoV-2 spike protein/ SARS-CoV-1 antibody, the closest in genomic sequence. As speculated, the virus interacts, at the molecular level, with either the IgM/IgG or ACE2, mainly through the hydrogen bonds formed by carboxyl and/or amino groups^[23–25]^, infrared spectroscopy (FTIR) should thus be the most revealing physical instrument^[6,26,27]^, to probe not only the bonding chemistry in a qualitative manner, but also the quantitative information, such as the number of the binding sites, leading to a direct measure of the virus attacking intensity. Employing FTIR, it was found by surprise that, the bonding numbers of the two antibody types are almost identical at lab/room temperature, or anything below 27°C. As a result, the antibodies were, astonishingly, rather nonspecific, resulting in a lower than 95% accuracy. On the other hand, such specificity to the SARS-CoV-2 antibody could be achieved at the human body temperature of 37°C, or at least beyond 31°C, when the quaternary protein structure would be further unfolded, to match that of the antibodies specifically, where the two antibody types reached corresponding numbers of available hydrogen bonds (19 vs 12) in their structures. Our results undoubtedly calls for the necessity of simulating human body temperature in the future antibody diagnosis, especially during the search for the possible vaccines, as the binding uniqueness, or the specificity of SARS-CoV-2 IgM/IgG, could only be fully obtainable at a warmer temperature, rather than under the usual lab/room temperature of 20°C - 25°C, where most of the vaccine research is conducted.

## Results & Discussion

Based on the nature of hydrogen bonding between the pairs of carboxyl and amino groups (**Fig. 1a**), samples of antibodies and spike proteins were mixed, and undergone FTIR spectroscopic examination conditioned by variable temperature (**Fig. 1b**). As the protein sample concentrations are quite low (0.1wt%), the attenuated total reflectance (ATR) type of FTIR has been adopted, which entertains a tiny sample amount of several microliters, 0.1% error margin in the absorption measurement, and 0.5cm^-1^ along the wavenumber scan. To facilitate the possible temperature variation, an electric heating element, and a thermometer were attached to the metallic plate of the ATR assembly, which also acts as an ideal heat sink (**Fig. 1b**). To include all the possible hydrogen bond stretching modes (**Fig. 1a**), the entire mid infrared spectrum was scanned for the two 1:1 mixtures of, SARS-CoV-2 spike protein/ SARS-CoV-2 antibody (S + Antibody 2), and SARS-CoV-2 spike protein/ SARS-CoV-1 antibody (S + Antibody 1) (**Fig. 1c, 1d**), where many absorption bands are observed which could be attributed to the proteins, such as: 2500-3500cm^-1^ from -OH, 2400-3200cm^-1^ from NH_2_, 1650cm^-1^ from the peptide bonds, and 800-1000 cm^-1^ from C-H^[6]^. However, due to the low concentration, they are mostly overlapped by the FTIR of the buffer solution PBS, which is dominated by water (0.1wt%NaCl +0.002wt%KCl +0.01wt%Na_2_HPO_4_ +0.002%wtKH_2_PO_4_ +99^+^wt%H_2_O), such as: 2500-3500cm^-1^ from -OH stretching, 2250-2500cm^-1^ from H-O-H bending, 1650 cm^-1^ from -OH scissoring, and 800-1000 cm^-1^ from H_2_O liberation, which could neither be easily resolved by D_2_O replacement,^[28,29]^ although absorption band at around 1630cm^-1^ does show some protein variation^[6]^. Fortunately, there exists a FTIR absorption band exclusively associated with the proteins, but not overshadowed by the solvent at 1550 cm^-1^ (**Fig. 1c-d, 2a-b**) which is caused by the carboxyl and amino groups in the amino acid^[30]^, allowing us to probe the protein bonding variation.

**Figure 1.**
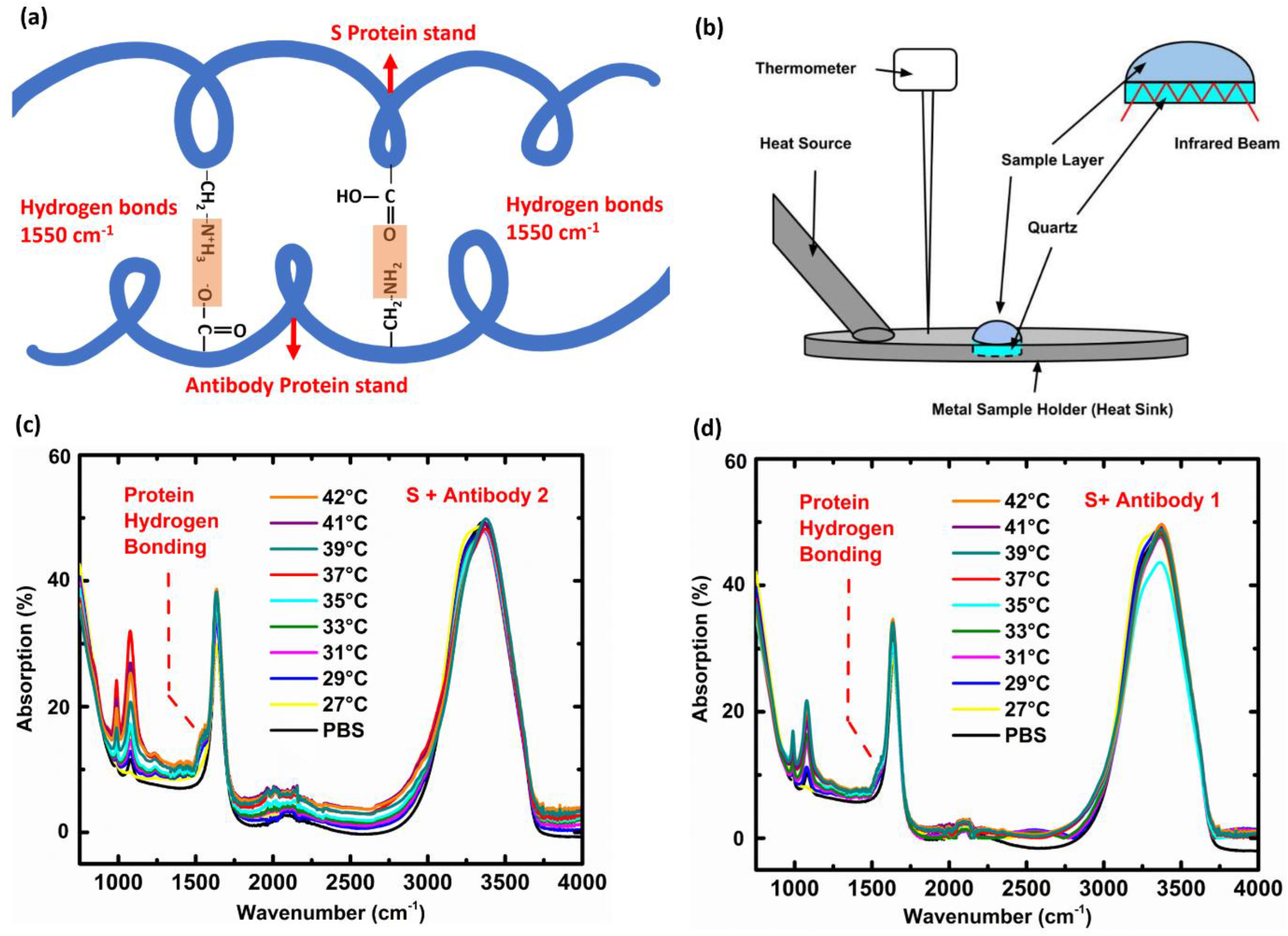
(a) Schematic illustration of the hydrogen bonding between carboxyl and amino groups of the protein strands, representing a typical intermolecular binding for the secondary structure of proteins. (b) The FTIR apparatus, the ATR model has been adopted to improve the signal noise ratio, which offers 0.1% error margin in the absorption measurement, and 0.5cm^-1^ along the wavenumber scan. To facilitate the possible temperature variation, an electric heating element, and a thermometer were attached to the metallic plate of the ATR assembly, which also acts as an ideal heat sink. Samples of antibodies and spike proteins were mixed together, and undergone FTIR spectroscopic examination conditioned by variable temperatures. (c) The FTIR scan of the sample mixture of SARS-CoV-2 spike protein/ SARS-CoV-2 antibody (S + Antibody 2) in PBS solution. Comparing the FTIR signal of the PBS solvent, it indicates that 1550 cm^-1^ was the only absorption band exclusively associated with the proteins, but not overshadowed by the PBS solvent. (d) The same FTIR scan of the sample mixture of SARS-CoV-2 spike protein/ SARS-CoV-1 antibody (S + Antibody 1) in PBS solution.

**Figure 2.**
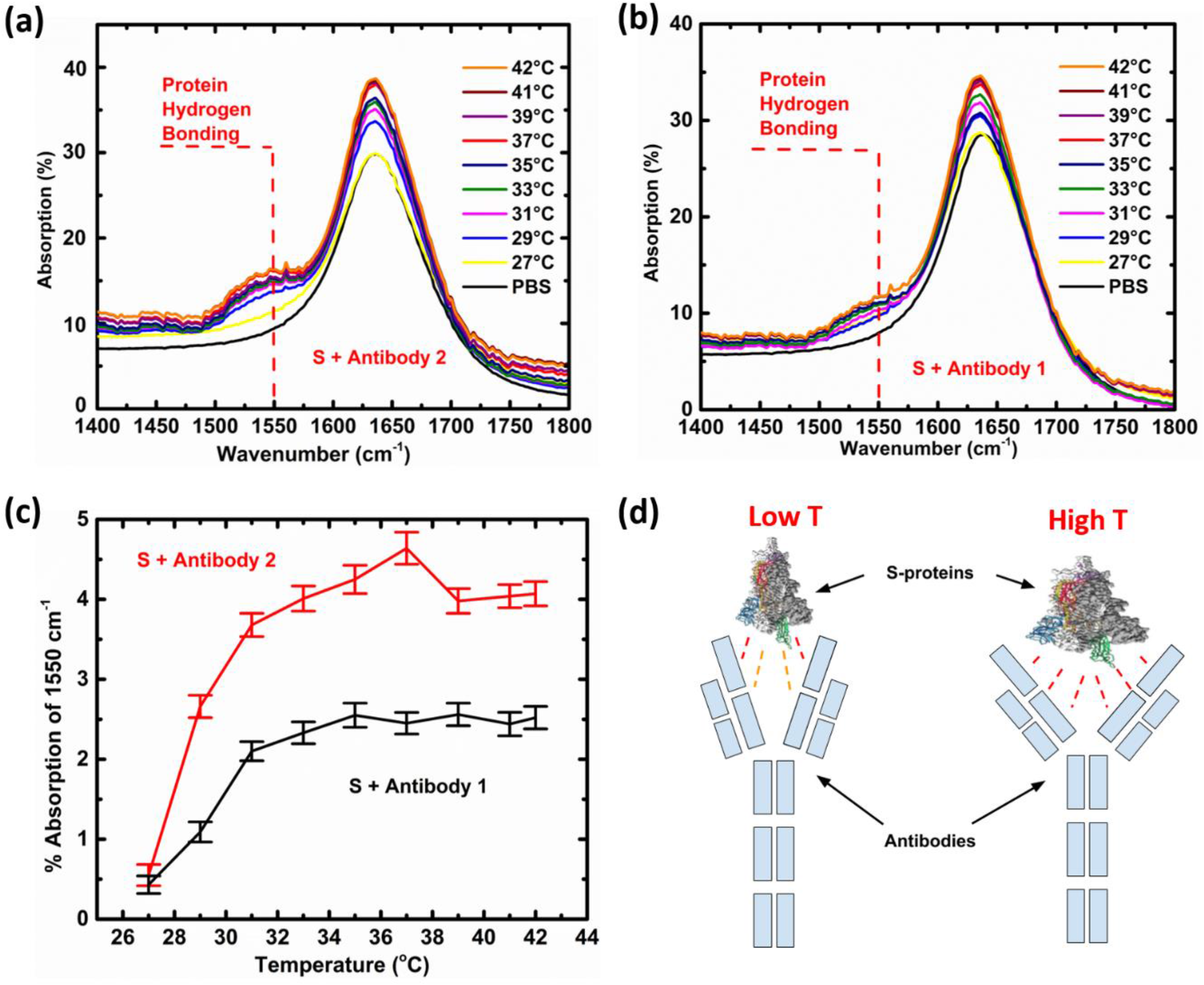
(a) Detailed FTIR scan of the sample mixture of SARS-CoV-2 spike protein/ SARS-CoV-2 antibody (S + Antibody 2) in PBS solution, within the wavenumber range of 1400-1800cm^-1^. (b) Detailed FTIR scan of the sample mixture of SARS-CoV-2 spike protein/ SARS-CoV-1 antibody (S + Antibody 1) in PBS solution, in the wavenumber range of 1400-1800cm^-1^. (c) The infrared absorptions of 1550cm^-1^ under variable temperature of the sample combination of SARS-CoV-2 spike protein/ SARS-CoV-2 antibody, versus the sample combination of SARS-CoV-2 spike protein/ SARS-CoV-1 antibody, obtained from the difference between the peak near 1550cm^-1^ and the trough near 1480cm^-1^, after subtracting that of the buffer solution. The associated error bars were estimated by combining the instrumental error, and the possible deviation involved in the computation. A strong temperature dependence is observed, where the bonding number of the antibody is enhanced sharply beyond 31°C, rather than at the usual room temperature. (d) The possible temperature influence on the structure of spike protein and antibodies, where only the top part of the IgM, similar to IgG, was depicted: a higher temperature increases the probability of protein quaternary structure unfolding, and exposes more binding sites, hence more bonds.

More details can be found from (**Fig. 2a,b**), where a strong temperature dependence is observed by surprise (**Fig. 2c**), for the absorption band of 1550cm^-1^, after the subtraction of the buffer solution absorption. At 27°C, for example, similar absorbance of 0.5% and 0.4% were found at 1550 cm^-1^, between the mixtures of the SARS-CoV-2 spike protein/SARS-CoV-2 antibody and the SARS-CoV-2 spike protein/ SARS-CoV-1 antibody, respectively. However, the absorbance increased to 4.4% and 2.5% at 37°C, respectively, presenting not only a significant amount of enhancement, but also a 72% discrepancy between the two types of antibodies. The enhanced absorption at 1550 cm^-1^ clearly indicates the augmentation of the hydrogen bonding numbers, rather than strengthening the binding itself, as the latter would relocate the band to a larger wavenumber, corresponding to the higher energy. Therefore, the result implies that, there are fewer bonds, or less specificity, between the spike protein and its antibody at lab/room temperature of below 27°C, when the S association with the SARS-CoV-2 antibody and SARS-CoV-1 antibody becomes similar, leading to the possible false conclusion. The difference or specificity will however be enhanced at higher temperatures, at least beyond 31°C, when the ratio becomes almost 19 vs 11, suppressing effectively the binding of the wrong types of antibody (**Fig. 2c**).

Such an increase in bonding numbers at elevated temperatures is likely caused by the swelling or unfolding of the protein strands (**Fig. 2d**), which can be further verified by **Fig. 3a, b, & c**, where the S proteins and antibodies were assessed by FTIR separately. The absorption band at 1550 cm^-1^ followed a similar trend in temperature (**Fig. 3d**), presumably attributed to the thermal agitation, when the likelihood increases for such binding sites to pair, either between the neighbouring proteins, or amongst various strands within a protein, or even along a protein strand (**Fig. 3e**). On the other hand, when the spike proteins and antibodies were mixed, the bond pairing occurs mainly across the different species (**Fig. 1a**), in order to obey the universal law of entropy maximization. The latter can be confirmed by the comparison of **Fig. 2c & Fig. 3d**, where the absorbance of the mixture is much larger than the sum of each individual component beyond 31°C, viz., when the signal would be the average of the two, should there be no mutual interactions, as the samples were all prepared by the same 0.1wt% in the solvent. In fact, the FTIR contribution caused by the antibody binding to the spikes could be extracted from the difference between the mixture absorption and the superposition of the individual. Beyond the most distinct band at 1550 cm^-1^, similar trends can also be observed in other absorption bands, such as the 980 cm^-1^ and 1080 cm^-1^, attributed to -C=O in serine and -OH in threonine^[6]^. As can be judged from **Fig. 1, 2&3**, both mixture absorptions were similar to the superposition of the individual absorptions at lower temperatures, indicating the relocation of the bond pairing from the individual proteins to the mixtures, whose binding number was relatively small. At higher temperatures, on the other hand, the absorption of both mixtures was much higher than the superposition of the individual absorption, representing a larger binding number. All this can be further verified by an extra absorption band found near 850 cm^-1^, due exclusively to the binding between the virus spike proteins and the antibodies, since it was absent in the individual spectrum of neither.

**Figure 3.**
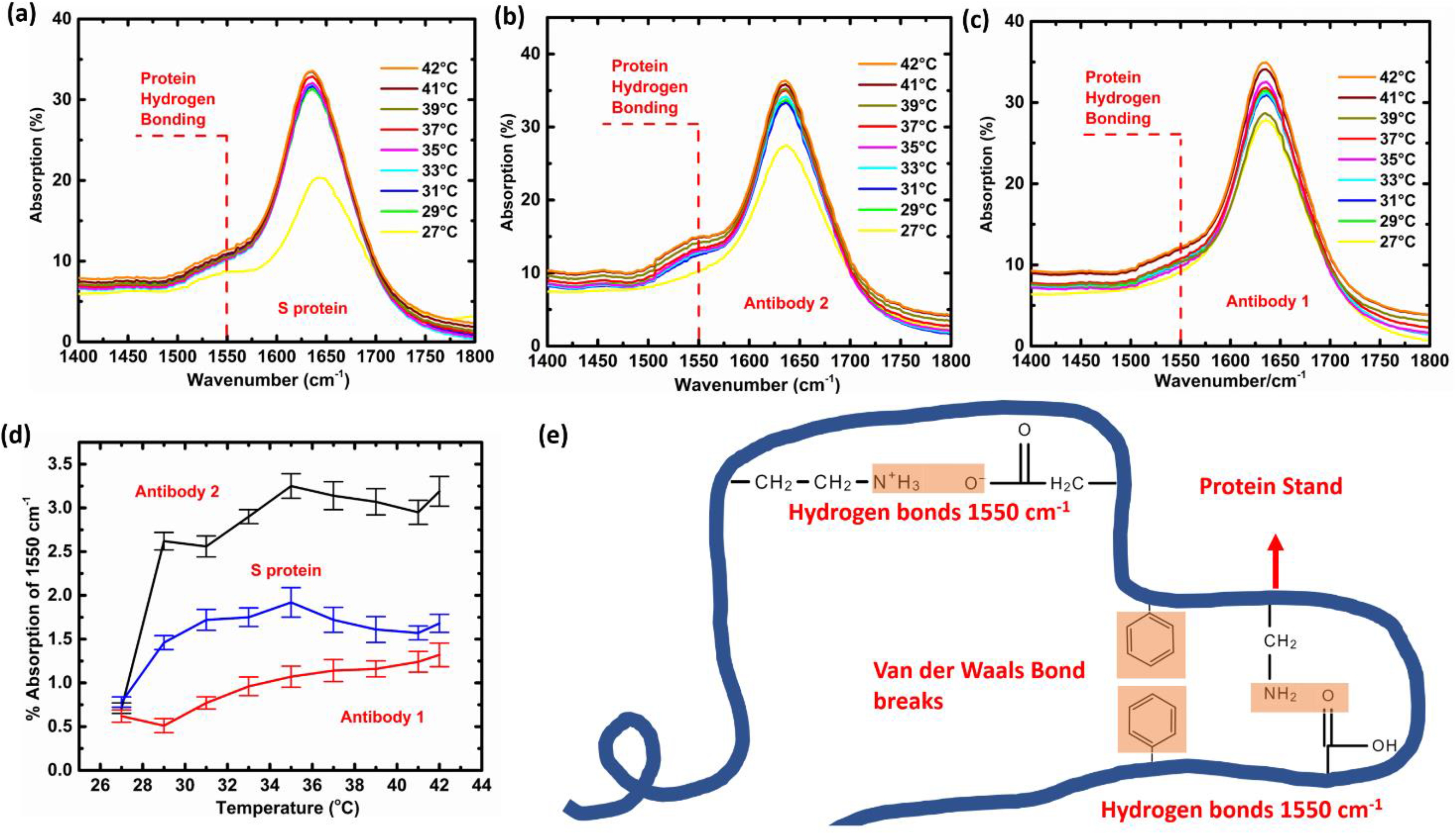
(a) The FTIR scan of SARS-CoV-2 spike protein (S protein) in PBS solution, for the wavenumber range of 1400-1800cm^-1^. (b) The FTIR scan of SARS-CoV-2 antibody (antibody 2) in PBS solution, for the wavenumber range of 1400-1800cm^-1^. (c) The FTIR scan of SARS-CoV-1 antibody (antibody 1) in PBS solution, for the wavenumber range of 1400-1800cm^-1^. (d) The infrared absorption of 1550cm^-1^ under variable temperature, of SARS-CoV-2 spike protein, SARS-CoV-2 antibody, and SARS-CoV-1 antibody, respectively, given by the difference between the crest near 1550cm^-1^, and the trough near 1480cm^-1^. The associated error bars were estimated by combining the instrumental error, and the possible deviation involved in the computation. All 3 proteins followed a similar trend in temperature, which is attributed to the thermal agitation, when the likelihood increases for such binding sites to pair, either between the neighbouring proteins, or amongst various strands within a protein, or even along a protein strand. (e) Schematic illustration of the hydrogen bonding between carboxyl and amino groups within a protein strand, showing the intra-molecular binding of the quaternary structure of proteins, when the Van der Waals bonds between large pendant groups, such as phenyl, were disrupted, which causes unfolding and exposing more bonding sites.

The gradual exposure of the bonding sites, caused by thermally enhanced unfolding of the quaternary protein structure, was further evidenced by the diminishing Van der Waals bonds in the FTIR of spike protein and antibodies, as shown by the 750 cm^-1^ band (**Fig. 4a, b, c**), which were obtained from the measurement of more concentrated samples via solvent evaporation, to achieve stronger absorption, as well as to reduce the influence of solvent. Being closer to the backbones, the opposite temperature dependence is anticipated for the phenyl groups to the amino/carboxyl groups. Rather than allowing for more hydrogen bonds by the temperature increase, the absorption band of 750 cm^-1^, gradually diminishes at elevated temperature (**Fig. 4d**), which indicates the collapse of Van der Waals bonds between the phenyl groups along the backbone. They must be thermally fractured, allowing the protein strands to further unfold, exposing more hydrogen bonding sites for pairing. The thermally activated unfolding promotes in turn the enhancement in the number of hydrogen bonds between the spike proteins and their antibodies. In the meantime, **Fig. 4d** also verifies the observation by comparing **Fig. 2c & Fig. 3d**, viz., the mixing produces more Van der Waals bonds, due to the entropy maximization, although it occurs at lower temperature for the phenyl groups, and does not differentiate the two antibody types. At higher temperature, this is further escalated between the S protein and the SARS-CoV-2 antibody, due to the hydrogen bonding, as shown in **Fig. 2c**.

**Figure 4.**
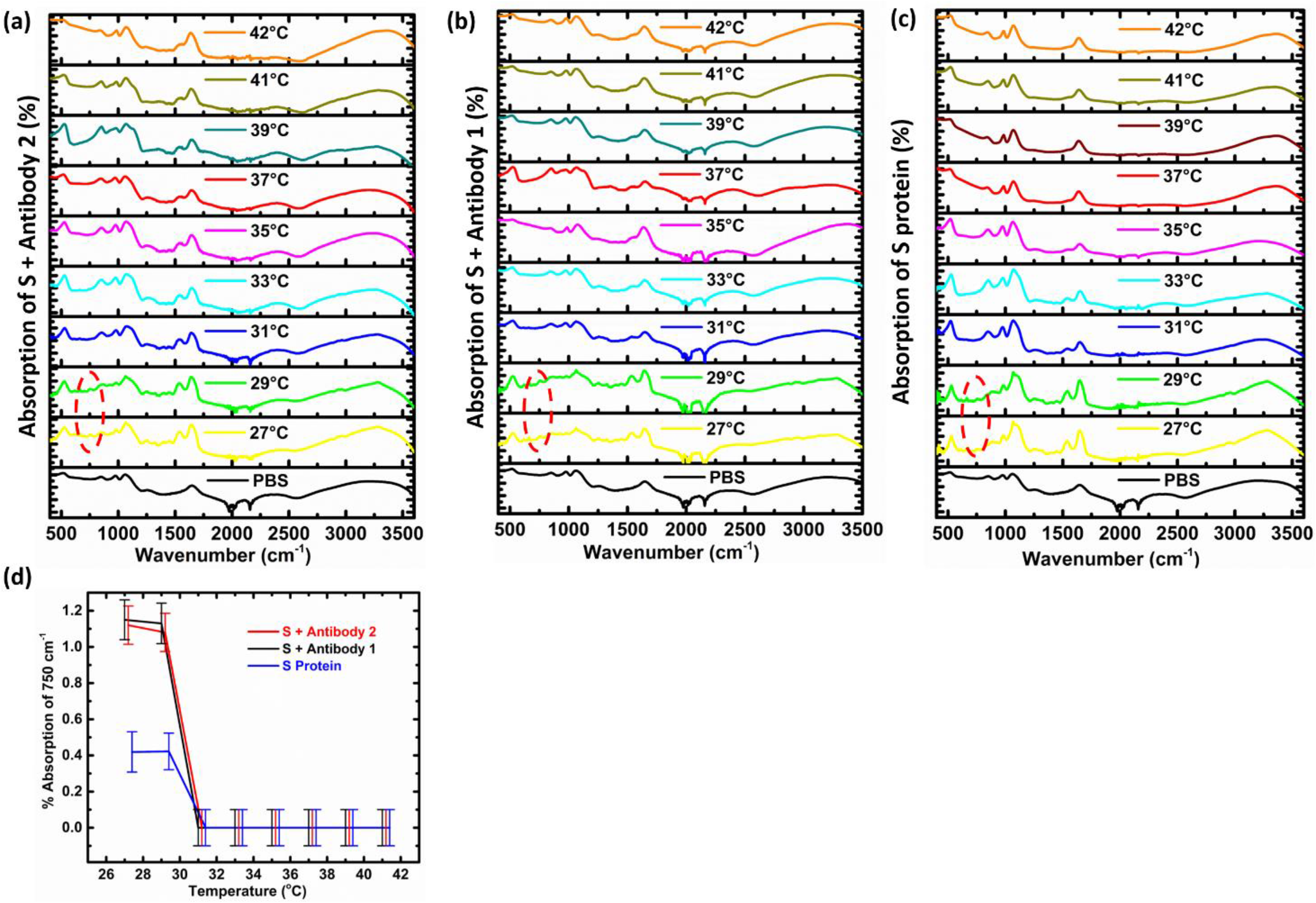
(a) The FTIR scan of SARS-CoV-2 spike protein/ SARS-CoV-2 antibody (S + Antibody 2), obtained from more concentrated samples, showing a visible 750 cm^-1^ bands at 27-29°C (red circle), exclusively associated with the proteins, but not found in the PBS solvent, attributed to the possible Van der Waals bonding between the large pendant group such as phenyl. (b) Similar FTIR result of SARS-CoV-2 spike protein/ SARS-CoV-1 antibody (S + Antibody 1) obtained from more concentrated samples, showing a visible 750 cm^-1^ bands at 27-29°C (red circle). (c) Similar FTIR result of SARS-CoV-2 spike protein alone, obtained from more concentrated samples, showing a visible 750 cm^-1^ bands at 27-29°C (red circle). (d) The infrared absorption of 750cm^-1^ under variable temperature of SARS-CoV-2 spike protein/ SARS-CoV-2 antibody (S + Antibody 2), SARS-CoV-2 spike protein/ SARS-CoV-1 antibody (S + Antibody 1), and the spike protein alone, obtained from the difference between the peak near 750cm^-1^ and the trough near 725cm^-1^, after subtracting that of the buffer solution. The associated error bars were estimated by combining the instrumental error, and the possible deviation involved in the computation.

As a matter of fact, similar evidence can also be traced from the recent literature, where the binding sites differ between SARS-CoV-2 antibody and SARS-CoV-1 antibody^[5,31–33]^. For example, amino acid sequencing was conducted for the receptor-binding domains (RBD) of SARS-CoV-2 and SARS-CoV-1 spikes. It was discovered that only 23% of the sequence mismatches between the two spikes, which calls for the further analysis of binding site numbers within this region^[19]^. From the literature, it was found that there are 19 -N l· groups and 0 -COOH group in the side chains of the SARS-CoV-2 spike RBD backbone, whereas their SARS-CoV-1 counterpart contains 11 -NH_2_ groups and 5 -COOH groups only^[19,34]^. As these groups form potential binding sites to the antibody, ideally the corresponding SARS-CoV-2 antibody should bear 19 -COOH groups, versus the SARS-CoV-1 antibody of only 11 -COOH groups and 5 -NH_2_ groups (**Table 1**). This is verified by a just published paper, which not only confirms the number of hydrogen bonds, but also supports the new picture brought together by our results^[33]^. As shown by **Fig. 5**, where the heavy chains of the HVR in the antibody 2 and 1 were compared, which involve 19 -COOH groups and 1 -NH_2_ group in antibody 2, and 12 -COOH groups and 4 -NH_2_ groups in antibody 1, rendering a very close match to our predication (**Table 1**). Although the implication of the slight mismatch remains to be further explored, at least the analysis well correlates quantitatively the FTIR results of 1550 cm^-1^ band in **Fig. 2c**, where the absorption peak of SARS-CoV-2 spike and SARS-CoV-2 antibody mixture is about 72% stronger than that of the SARS-CoV-2 spike and SARS-CoV-1 antibody, coincide with the ratio of 19 vs 11, when the proteins are equilibrated at 37°C.

**Figure 5.**
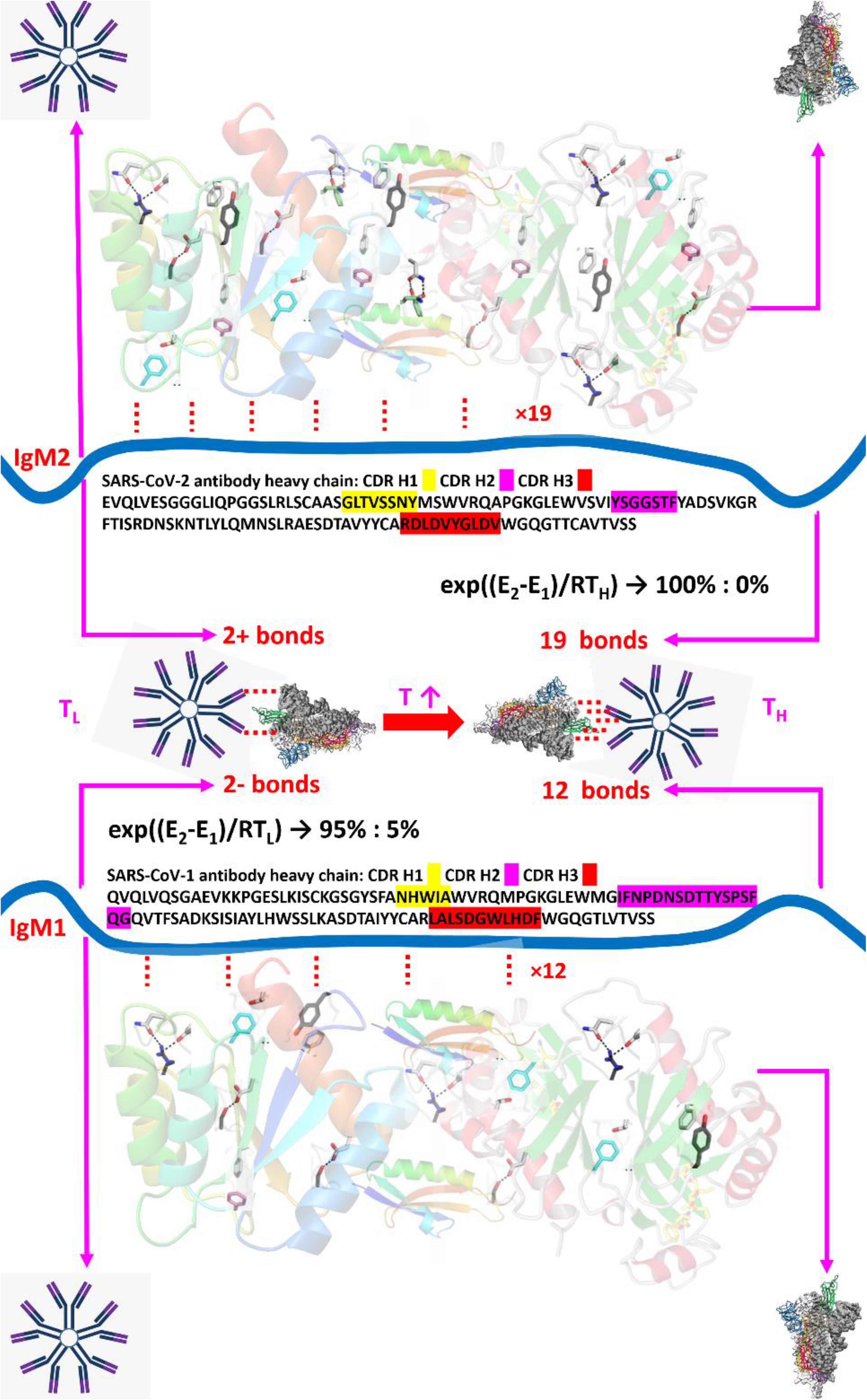
Comparison of antibody 2 and antibody 1 amino acid sequence with respect to the binding difference to the S protein. The IgM were depicted by the blue ribbon in the middle, and the S proteins represented by the screw ribbons in the top and bottom, where some of representative epitope residues in the S protein were labelled. The sequence of the antibodies were obtained from the literature^[32,33]^. Both antibodies form only 2+ hydrogen bonds at room temperature due to the folding of protein structure, while such number would increase to 19 and 12, respectively, at human body temperature. As can be calculated by the Arrhenius equation, around 5% of the S protein would bind with antibody 1 at room temperature, resulting in unavoidable testing inaccuracy which could only be eliminated at body temperature.

**Table 1.**
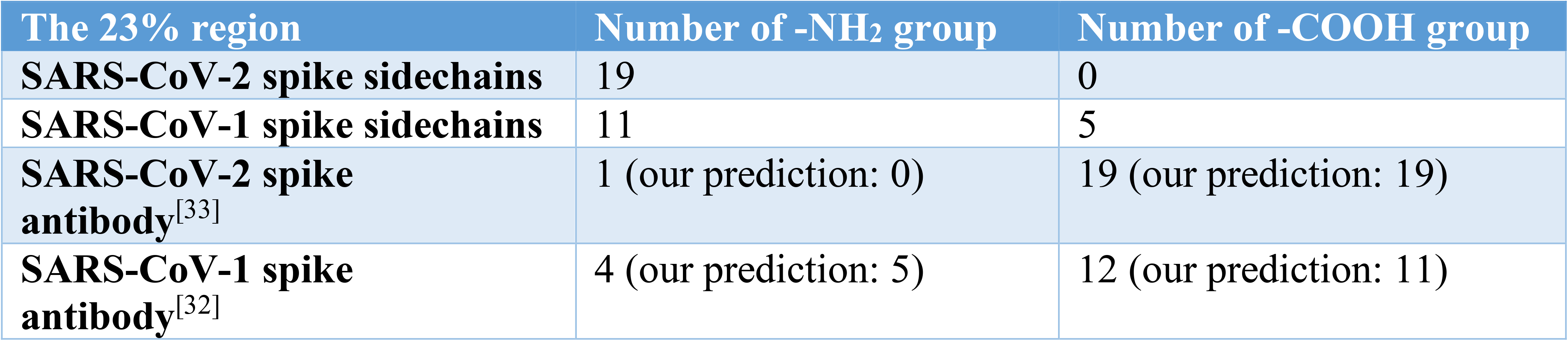
The number of binding sites in SARS-CoV-2/SARS-CoV-1 spike proteins and their antibodies^[19,34]^.

| The 23% region | Number of -NH<sub>2</sub> group | Number of -COOH group |
| --- | --- | --- |
| SARS-CoV-2 spike sidechains | 19 | 0 |
| SARS-CoV-1 spike sidechains | 11 | 5 |
| SARS-CoV-2 spike antibody<sup>[33]</sup> | 1 (our prediction: 0) | 19 (our prediction: 19) |
| SARS-CoV-1 spike antibody<sup>[32]</sup> | 4 (our prediction: 5) | 12 (our prediction: 11) |

Exploring further the consequence of the temperature influence on the bonding between virus and antibody, vis-à-vis the gradual unfolding of the protein structure, quantitative information may be obtained through Arrhenius equation, where the probability of bond breaking is related to an exponential function of the bonding energy E over the temperature T: exp(-E/RT), where R is the gas constant. Such equation predicts the stability of the protein and the unfolding of the quaternary structures. For example, the possibility to break a hydrogen bond of 19kJ/mol on the secondary structure, is almost negligible (0.04% at 295K - 0.06% at 310K), whereas a quaternary structure of 1.0-5.0kJ/mol, will much more likely be affected (15%-68%) by the same temperature change, when more bonding sites on the spike RBD area are exposed, resulting in tighter binding to the antibody. Finally, following thermodynamics, if we designate the bond energy, times the number of such bonds per mole for the two antibody types, by E_1_ and E_2_ (kJ/mol), the ratio of bonded antibody numbers, [Ab_2_]/[Ab_1_], will then be given by exp((E_2_ - E_1_)/RT), rather than the layman’s guess of E_2_/E_1_. Following the result of **Fig. 2c**, where the 0.1% absorbance discrepancy of the two antibodies at 27°C corresponds to an energy difference of 7.5 KJ/mol, the antibody number ratio can thus be calculated by the exponential to become 20:1, give rise to 5% of the test uncertainty.

## Conclusion

In conclusion, we have further probed interactions between SARS-CoV-2 spike protein and SARS-CoV-2 versus the closest kin of SARS-CoV-1 antibodies. Beyond the usual genomic sequencing, infrared spectroscopic measurement allows us to probe not only the presence of the hydrogen bonds between spike protein and antibodies, but also the disparity in the number of bonding sites among the antibodies, given by the thermodynamic exponential ratio. The binding strength between SARS-CoV-2 spike and SARS-CoV-2/SARS-CoV-1 antibodies was found to be temperature dependent, where a higher temperature raises the probability of protein quaternary structure unfolding, which exposes more binding sites, hence more bond numbers. This was also confirmed by the similar observations for the proteins themselves, especially from the diminishing of Van der Waals bonds. In the meantime, the relative absorbance matches at 37°C the hydrogen bonding numbers of the two antibody types (19 vs 11), whereas the ratio of bonds at 27°C, calculated by thermodynamic exponentials rather than by the layman’s guess, is about 20:1, leading to the 95% accuracy at its best. Moreover, the virus/antibody bonding specificity is enhanced at human body temperature, instead of room temperature, due to the possible protein unfolding by thermal agitation, leading to the necessity of conducting the vaccine research at 37°C to identify various viruses. The temperature variation and the bond numbers not only establish a linkage between the pandemic and molecular level query^[35,36]^, but also provide a new dimension in raising the accuracy of the ongoing Rapid Antibody Test, which will be beneficial to the SARS-CoV-2 vaccine exploration and virus research in general.

## Acknowledgement

Thanks are due to Beijing Institute of Technology for the FTIR support, Prof. L. Zhang of University of Toronto Health Network for comments, and J. McLellan of University of Texas at Austin, for the artwork of SARS-CoV-2 Spike proteins. Also acknowledged is the NSERC of Canada.

## Author contributions

RTW mastered every stage of the research, AFX performed measurements, KJX helped manuscript preparation, TS supported measurement, GX initiated and supervised the research.

## Experimental

SARS-CoV-2 Spike RBD-His (V367F) Recombinant Protein (Purity: > 95 % as determined by SDS-Page, protein concentration: 0.1wt%), SARS-CoV-2 antibody (antibody 2, also named SARS-CoV-2 (2019-nCoV) Spike Neutralizing Antibody), and SARS-CoV-1 antibody (antibody 1, also named SARS-CoV/SARS-CoV-2 Spike S1 antibody) (Purity: > 95 % as determined by SDS-Page, protein concentration: 0.1wt%) were all purchased from Sino Biological Inc. PBS solution (phosphate buffered solution containing 0.1wt%NaCl +0.002wt%KCl +0.01wt%Na_2_HPO_4_ +0.002wt%KH_2_PO_4_ +99^+^wt%H_2_O) was purchased from Solarbio Corp.

A Nicolet iS50 FT-IR spectrometer was adopted to acquire the FTIR spectra, in wavenumber range of 500-4000 cm^-1^ by the attenuated total reflectance (ATR) mode. Samples were placed in a sample stage made of quarts. The specified wavenumber precision is 0.5 cm^-1^ and that of the absorbance is 0.1%. A blank run was first conducted to ensure the calibration. An electric heating element was installed to change the temperature of the sample stage. The temperature was recorded via electronic thermometer with an accuracy of ± 0.1°C. The liquid samples were pipetted to ensure the same amount of 3μl each time.

The acquired FTIR data were inverted to obtain the %absorption from (100% - transmittance %). PBS solvent contributions can be eliminated by subtracting the spectra of PBS under identical conditions. Infrared absorption at 750cm^-1^ and 1550cm^-1^ were determined by the difference between the crest and the trough. The associated error bars were estimated by combining the instrumental error, and the possible deviation involved in the computation.

## Reference

[1] L. Guo, L. Ren, S. Yang, M. Xiao, D. Chang, F. Yang, C. S. Dela Cruz, Y. Wang, C. Wu, Y. Xiao, L. Zhang, L. Han, S. Dang, Y. Xu, Q. Yang, S. Xu, H. Zhu, Y. Xu, Q. Jin, L. Sharma, L. Wang, J. Wang, Clin. Infect. Dis. 2020, DOI:10.1093/cid/ciaa310.

[2] Y. W. Tang, J. E. Schmitz, D. H. Persing, C. W. Stratton, J. Clin. Microbiol. 2020, 58, DOI:10.1128/JCM.00512-20.

[3] X. Tian, C. Li, A. Huang, S. Xia, S. Lu, Z. Shi, L. Lu, S. Jiang, Z. Yang, Y. Wu, T. Ying, Emerg. Microbes Infect. 2020, 9, 382.

[4] Z. Li, Y. Yi, X. Luo, N. Xiong, Y. Liu, S. Li, R. Sun, Y. Wang, B. Hu, W. Chen, Y. Zhang, J. Wang, B. Huang, Y. Lin, J. Yang, W. Cai, X. Wang, J. Cheng, Z. Chen, K. Sun, W. Pan, Z. Zhan, L. Chen, F. Ye, J. Med. Virol. 2020, DOI:10.1002/jmv.25727.

[5] Q. Wang, Y. Zhang, L. Wu, S. Niu, C. Song, Z. Zhang, G. Lu, C. Qiao, Y. Hu, K. Y. Yuen, Q. Wang, H. Zhou, J. Yan, J. Qi, Cell 2020, 181, 894.

[6] A. Barth, Biochim. Biophys. Acta - Bioenerg. 2007, 1767, 1073.

[7] N. H. Andersen, J. Am. Chem. Soc. 2001, 123, 12933.

[8] B. S. Graham, Science (80-.). 2020, 368, 945.

[9] P. J. Hotez, D. B. Corry, M. E. Bottazzi, Nat. Rev. Immunol. 2020, 20, 347.

[10] A. Petherick, Lancet (London, England) 2020, 395, 1101.

[11] S. Jiang, C. Hillyer, L. Du, Trends Immunol. 2020, 41, 355.

[12] F. Amanat, F. Krammer, Immunity 2020,.

[13] E. Padron-Regalado, Infect. Dis. Ther. 2020, DOI:10.1007/s40121-020-00300-x.

[14] K. V. Iserson, Cambridge Q. Healthc. Ethics 2020, DOI:10.1017/s096318012000047x.

[15] M. Infantino, V. Grossi, B. Lari, R. Bambi, A. Perri, M. Manneschi, G. Terenzi, I. Liotti, G. Ciotta, C. Taddei, M. Benucci, P. Casprini, F. Veneziani, S. Fabbri, A. Pompetti, M. Manfredi, J. Med. Virol. 2020, DOI:10.1002/jmv.25932.

[16] R. S. Khan, I. U. Rehman, Expert Rev. Mol. Diagn. 2020, DOI:10.1080/14737159.2020.1766968.

[17] G. Peng, Y. Yang, J. R. Pasquarella, L. Xu, Z. Qian, K. V. Holmes, F. Li, J. Biol. Chem. 2017, 292, 2174.

[18] M. Hoffmann, H. Kleine-Weber, S. Schroeder, N. Krüger, T. Herrler, S. Erichsen, T. S. Schiergens, G. Herrler, N. H. Wu, A. Nitsche, M. A. Müller, C. Drosten, S. Pöhlmann, Cell 2020, 181, 271.

[19] W. Tai, L. He, X. Zhang, J. Pu, D. Voronin, S. Jiang, Y. Zhou, L. Du, Cell. Mol. Immunol. 2020, 17, 613.

[20] W. D., W. N., C. K.S., G. J.A., H. C.-L., A. O., G. B.S., D. Wrapp, N. Wang, K. S. Corbett, J. A. Goldsmith, C.-L. Hsieh, O. Abiona, B. S. Graham, J. S. McLellan, Science (80-.). 2020, 367, 1260.

[21] F. Wu, S. Zhao, B. Yu, Y. M. Chen, W. Wang, Z. G. Song, Y. Hu, Z. W. Tao, J. H. Tian, Y. Y. Pei, M. L. Yuan, Y. L. Zhang, F. H. Dai, Y. Liu, Q. M. Wang, J. J. Zheng, L. Xu, E. C. Holmes, Y. Z. Zhang, Nature 2020, 579, 265.

[22] Y. Gao, L. Yan, Y. Huang, F. Liu, Y. Zhao, L. Cao, T. Wang, Q. Sun, Z. Ming, L. Zhang, J. Ge, L. Zheng, Y. Zhang, H. Wang, Y. Zhu, C. Zhu, T. Hu, T. Hua, B. Zhang, X. Yang, J. Li, H. Yang, Z. Liu, W. Xu, L. W. Guddat, Q. Wang, Z. Lou, Z. Rao, Science 2020, DOI:10.1126/science.abb7498.

[23] D. C. Ekiert, F. Wang, I. A. Wilson, P. G. Schultz, V. V. Smider, Biophys. J. 2014, 106, 438a.

[24] M. N. Nguyen, M. R. Pradhan, C. Verma, P. Zhong, Bioinformatics 2017, 33, 2971.

[25] R. A. Copeland, in Enzymes, 2003, pp. 11–41.

[26] K. Murayama, M. Tomida, Biochemistry 2004, 43, 11526.

[27] I. H. M. van Stokkum, H. Linsdell, J. M. Hadden, P. I. Haris, D. Chapman, M. Bloemendal, Biochemistry 1995, 34, 10508.

[28] S. Hume, G. Hithell, G. M. Greetham, P. M. Donaldson, M. Towrie, A. W. Parker, M. J. Baker, N. T. Hunt, Chem. Sci. 2019, 10, 6448.

[29] N. Kitadai, T. Sawai, R. Tonoue, S. Nakashima, M. Katsura, K. Fukushi, J. Solution Chem. 2014, 43, 1055.

[30] M. Nara, M. Tasumi, M. Tanokura, T. Hiraoki, M. Yazawa, A. Tsutsumi, FEBS Lett. 1994, 349, 84.

[31] M. Yuan, N. C. Wu, X. Zhu, C. C. D. Lee, R. T. Y. So, H. Lv, C. K. P. Mok, I. A. Wilson, Science (80-.). 2020, 368, 630.

[32] J. Liu, H. Shao, Y. Tao, B. Yang, L. Qian, X. Yang, B. Cao, G. Hu, H. Tachibana, X. Cheng, Clin. Vaccine Immunol. 2006, 13, 594.

[33] M. Yuan, H. Liu, N. C. Wu, C.-C. D. Lee, X. Zhu, F. Zhao, D. Huang, W. Yu, Y. Hua, H. Tien, T. F. Rogers, E. Landais, D. Sok, J. G. Jardine, D. R. Burton, I. A. Wilson, bioRxiv 2020, 2020.06.08.141267.

[34] W.-H. Chen, P. J. Hotez, M. E. Bottazzi, Hum. Vaccin. Immunother. 2020, 1.

[35] M. B. Araujo, B. Naimi, medRxiv 2020, 2020.03.12.20034728.

[36] R. A. Neher, R. Dyrdak, V. Druelle, E. B. Hodcroft, J. Albert, Swiss Med. Wkly. 2020, 150, w20224.

